# *Caenorhabditis elegans* model for admixed proteinopathy of Huntington’s and Alzheimer-type

**DOI:** 10.1101/648923

**Authors:** Wadim J. Kapulkin

## Abstract

Formation of proteinaceous deposits composed of abnormally aggregated proteins characterizes a range of pathological conditions. Proteinaceous inclusions detected in the neurodegenerative conditions such Huntington’s chorea (HD) and Alzheimer’s disease (AD) are often referred as pathognomic and regarded as causally implicated. Despite of differences in aetiology and underlying genetics, rare cases of combined HD and AD were reported and described as admixed proteinopathies. Mixed proteopathies are characterized by the co-occurrence of at least two types of abnormal aggregation-prone variants; pathological deposition of proteinaceous inclusions might however, affect different cell populations. Here, combining plaques derived from human ß-amyloid with mutant HTT-like polyglutamine inclusions in a cell-autonomous manner, we report on the *Caenorhabditis elegans* model for admixed proteinopathy of Alzheimer’s and Huntington’s type. We show both types of intracellular foci are formed *in vivo*: non-amyloidic extended polyglutamine derived inclusions and distinguish those from the presence of mature ß- amyloid fibers. We found that polyglutamines expanded above pathogenic threshold and ß-amyloid act synergistically to promote the progression of proteotoxicity in a temperature dependent manner. We further, implicate the *hsp-1* (the predominant *C. elegans* chaperone interacting with ER-routed Aß42) modulate the proteotoxic insults observed in combined proteopathy model. Our results demonstrate how the *in vivo* model of admixed proteopathy could be utilized to probe for human pathogenic variants confined to the same cellular type. In that perspective expanded aggregation-prone polyglutamines appended with fluorescent proteins could be regarded as ‘pathogenic probes’ useful in the proteotoxicity assays *i.e.* involving ß-amyloid and possibly other comparable models of disease-associated aggregation prone variants.We anticipate models of combined proteopathies will be informative regarding the underlying pathogenesis and provide the sensitized background for sophisticated screening. For example the combinatorial effects of multiple pathogenic aggregation-prone variants could be tested against mutant backgrounds and pharmacological compounds. Furthermore, we surmise the postulated synergistic actions might explain some of the overlaps in observed progression of clinical symptoms in HD and AD. We also formulate the conjecture regarding the polyQ containing proteins as a contributing factor in degenerative conditions associated with increased ß-amyloid formation and deposition.

## INTRODUCTION

Formation of proteinaceous inclusions occurs in a range of morbid conditions (Sipe et al. 2016). Pathological deposition of proteinaceous inclusions (often regarded as pathognomic in classical pathology) indicate for degenerative processes which may affect many cells types across tissues and organs. In particular, histopathological staining of congophilic material (inclusions reactive with Congo Red), is consistent with the deposition of abnormally aggregated proteins that share the structural properties common to amyloids (Tanskanen 2013). While amyloidosis can be observed in various aetiologically unrelated conditions in many animal species, the amyloidal nature of congophilic material detected in an aging brain is regarded as a hallmark of Alzheimer’s disease.

Alzheimer’s disease is classified as ‘conformational disease’ together with other age-related dementias and encephalopathies including Huntington’s chorea (Carrell &Lomas 1997). Referred broadly as proteopathies, conformational diseases, are characterized by the accumulation of aggregates composed of malfolded proteins (Kopito &Ron 2000). Accumulation of abnormally aggregated malfolded proteins set up the tissue-specific response of the chaperone system – the concept otherwise known as proteotoxicity (Ron 2002). Increased demand for cellular chaperoning in response to a formation of abnormal proteinaceous foci appears as a common component of the pathology underlying the neurodegenerative diseases (Wyttenbach &Arrigo 2013). While the chaperone response appears as a consistent feature associated with the degenerative process involving the deposition of aberrant proteins, the overall mechanisms and pathways underlying the disease, including the amyloidosis, remain poorly understood.

Given the enormous complexity of the vertebrate nervous system as well as general inaccessibility of the brain for clinical sampling purposes, *C. elegans* appears as an attractive model for the pathogenesis of degenerative diseases. *C. elegans* cells engineered to express pathogenic variants implicated in the pathogenesis of the neurodegenerative diseases, mount the robust chaperone response to abnormally aggregated proteins (Fonte et al. 2002; Link et al. 2003) and recapitulate the key features of the progression of cell degeneration *i.e.* amyloidosis (Link 1995; Link et al. 2001). Here, we report on the *C. elegans* model for admixed proteinopathy of Alzheimer’s and Huntington’s type. Both Alzheimer’s disease and Huntington’s chorea are associated with neurodegenerative features indicative for aberrant protein deposition observed in areas of affected brain (Spires &Hannan 2007). Despite the above parallels, the population of affected neuronal cells, particularly in prodromal and incipient stages is very different in both diseases (Braak &Braak 1991; Vonsattel &DiFiglia 1998; reviewed in Stroo et al. 2017) reflecting to the differences in underlying genetics, exact pathology sites and the disease onset (Wenk 2003; Walker 2007). The cases of combined proteinopathy of Alzheimer’s and Huntington’s type, are rare (Warner et al. 1994); Jellinger (1988) and others (references below) described Alzheimer’s-type lesions in Huntington’s disease. Here we utilized the genetic model organism to confine plaques derived from human ß-amyloid with mutant HTT-like polyglutamine inclusions in cell-autonomous manner.

## MATERIALS AND METHODS

### *C. elegans* strains and maintenance

Strains were maintained on NGM agar plates seeded with *E.coli* according to standard methods (Wood 1988). Classic genetic methods (Brenner 1974) were applied to construct double-transgenic lines. Briefly, transgenic hermaphrodite lines homozygous for mutant Huntigton-type polyglutamine appended with YFP {driven by unc-54 promoter (*unc-54p* from pPD30.38) Q0 AM134 (*rmIs126*[*unc-54p::Q0::yfp*] *X*); Q24 AM138 (*rmIs130*[*unc-54p::Q24::yfp*] *II*); Q35 AM140 (*rmIs132*[*unc-54p::Q35::yfp*] *I*); Q37 AM470 (*rmIs225*[*unc-54p::Q37::yfp*] *II*); and Q40 AM141 (*rmIs133*[*unc-54p::Q40::yfp*] *X*) genotypes referenced in (Morley et al. 2002; Brignull et al. 2006; Beam et al. 2012) developed and provided by the Morimoto lab (Northwestern University)} were mated with *him-6* males [*him-6*(*e1104*) CB1138 (Hodgkin et al. 1979)]. YFP^+^ *F1* males carrying above transgenes were crossed onto *unc-54p* driven ß-amyloid transgene (*dvIs2 II*) CL2006 (Link 1995) hermaphrodites to establish doubly-transgenic *F2* heterozygotes. Double-transgenic *F2* heterozygotes were selfed to assort double-homozygotes of desired genotypes (*dvIs2*^+/+^*; polyQ::yfp*^+/+^ respectively). Two stable doubly-homozygous isogenic lines of each YFP appended polyglutamine variant (derived from independent mating groups) were maintained for analysis while ensured that *him-6* allele segregated away.

### dsRNA mediated interference

*hsp-1* (IV) F26D10.3 dsRNA expressing construct (Kamath et al. 2003) were fed into double-transgenic lines, according to established protocols as described previously (Kapulkin et al. 2005, with relevant references). The efficacy of functional withdrawal of HSP70A was controlled by feeding individually in parallel with *bli-1* (causing fluid-filled blisters of the cuticle) and *unc-22* (causing distinct involuntary myoclonic twitches) RNAi constructs. Plasmids encoding dsRNA expressing constructs received from Silencing Genomes resource at the Cold Spring Harbor Laboratory (http://www.silencinggenomes.org/). dsRNA was administrated via HT115(DE3) *E.coli* [*F ^-^; RNase III* deficient*; T7 polymerase^+^*] strain used as an RNAi feeding vector as originally described in Timmons et al. (2001).

### Amyloid X-34 imaging

Transgenic hermaphrodites were stained *in vivo* for amyloid-plaques according to established protocols (Link et al. 2001) using X-34, a fluorescent derivative of Congo Red (Styren et al. 2000) kindy provided by William E. Klunk (University of Pittsburgh Department of Psychiatry - UPMC). Differentially illuminated fluorescent sections of X-34 stained double-homozygous transgenic hermaphrodites expressing ß-amyloid and polyglutamines appended with YFP were captured with MZ1 LSM 780 (Zeiss) laser scanning confocal microscope at MPI-CBG Light Microscopy Facility. Corresponding confocal sections were aligned and digitally merged in Fiji/ImageJ (Schindelin et al. 2012).

## RESULTS

We used extensively characterized C. elegans strains expressing human ß-amyloid peptide (Link 1995) and extended polyglutamines (Satyal et al. 2000) appended with YFP, to construct stable double-homozygotes expressing both transgenes in the same type of the cells. Stable double-homozygotes expressing both transgenes were assorted by selfing of double-heterozygotes derived from a cross of CL2006 (*dvIs2 II)* hermaphrodites mated with males carrying integrated arrays introducing the expanded polyglutamine stretch appended with YFP. Attempted crosses assorted independently double-homozygous lines co-expressing human ß-amyloid and expanded polyglutamines of two different lengths: expanded polyQ35 and polyQ40, appended with YFP, as well as polyQ non-extended control YFP. Attempted crosses failed to immediately assort double-homozygous lines in case of polyQ24-YFP and polyQ37-YFP due to disfavourable repulsion linkages of transgenic arrays integrated on chromosome II.

Representative individuals of stable doubly-homozygous lines expressing human ß-amyloid (*dvIs2 II)* and YFP appended to expanded polyglutamine stretches of two different lengths: polyQ35 (*rmIs132 I*) and polyQ40 (*rmIs133 X*) are shown in Fig. 1. The comparison of examined individuals of the above stable double transgenic lines reveals the difference in the general appearance: hermaphrodites expressing ß-amyloid along with polyglutamine Q40-YFP growing slower and reaching the smaller size (total body length reduced by ca ∼30%) when compared with animals co-expressing ß-amyloid along with polyQ35-YFP. Moreover, above differences in the growth rate affected by the length of expanded polyQ-YFP spanning the pathogenic threshold (Morley et al. 2002) appears to coincide with the overall movement deficit observed in the doubly-homozygous lines. The movement of doubly-homozygous lines expressing ß-amyloid along with Q35-YFP appears comparable to control non-extended YFP (*dvIs2; rmIs126*) line and singly transgenic parental (*dvIs2*) line CL2006 (a phenotype consistent with the presence of dominant *rol-6d(su1006)* used as a marker; not shown). In contrast, the movement of doubly-homozygous lines expressing ß-amyloid along with Q40-YFP, appears impaired in comparison with Q35-YFP, resulting in the *unc* (<u>unc</u>oordinated) phenotype imposed on the roller (*rol-6d*) movement (Fig. 2; and supplement movies: SupplMov.1 & SupplMov.2, <https://figshare.com> respectively).

**Figure 1.**
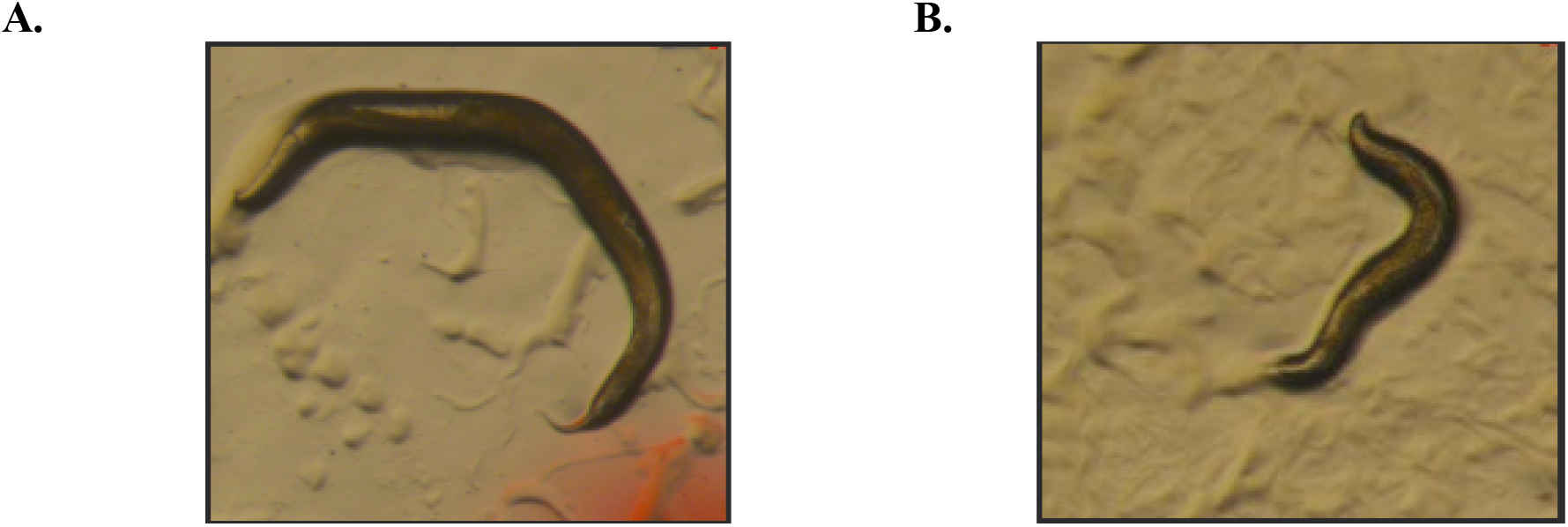
Representative individuals of stable doubly-homozygous lines expressing ß-amyloid (*dvIs2 II)* and YFP appended to expanded polyglutamine stretches of two different lengths: polyQ35 (*rmIs132 I*) and polyQ40 (*rmIs133 X*). A. Double-homozygote encoding for ß-amyloid and polyQ35-YFP (*dvIs2* ^+/+^; *rmIs132* ^+/+^) transgenes. Adult mobile roller hermaphrodite shown. Body posture typical for movement of *rol-6 (su1006)*, consistent with the presence of dominant roller marker derived from *dvIs2* amyloid-plaques encoding transgene. B. Atypical (*dvIs2* derived) roller morphology resulting from combined proteopathy composed of ß-amyloid and expanded polyQ40. Compare panel A (left) with immobile double-homozygote encoding for ß-amyloid and polyQ40- YFP (*dvIs2* ^+/+;^ *rmIs133* ^+/+^) presented on panel B (right). Age-matched young adults (grown for 4 days at 20°C) shown at the same magnification, as presented in supplement movies (SupplMov.1 and SupplMov.2, respectively).

**Figure 2.**
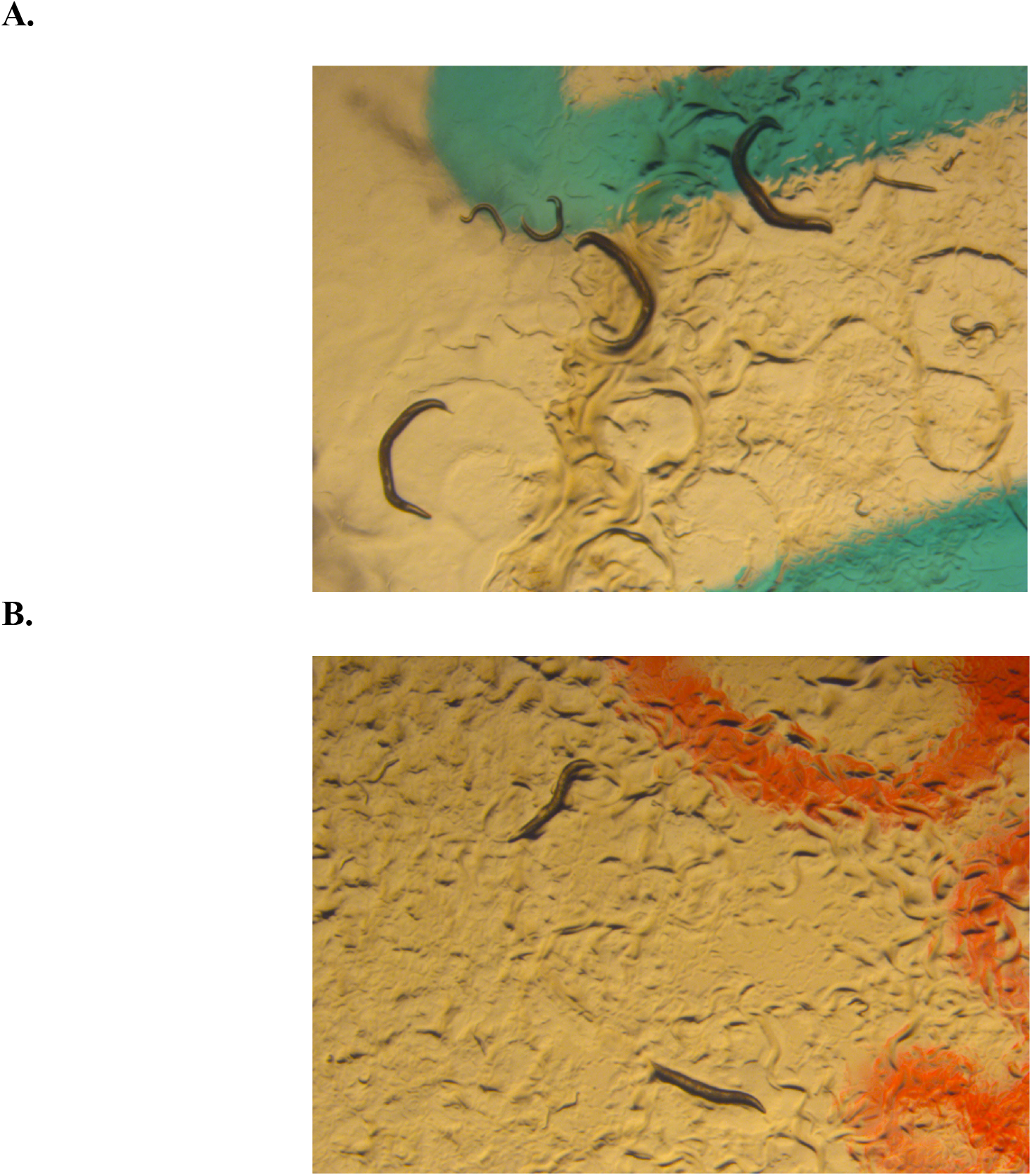
Plate appearance of stable doubly-homozygous lines expressing ß-amyloid (*dvIs2 II)* and YFP appended to expanded polyglutamine stretches of two different lengths: polyQ35 (*rmIs132 I*) and polyQ40 (*rmIs133 X*). A. Isogenic double-homozygotes encoding for ß-amyloid and polyQ35- YFP (*dvIs2* ^+/+^; *rmIs132* ^+/+^) transgenes. Three mobile adult roller hermaphrodites visible in addition to advanced larval stages representing the progeny. B. Atypical roller morphology resulting from combined proteopathy consisting of transgenic ß-amyloid and expanded polyQ40-YFP (*dvIs2* ^+/+;^ *rmIs133* ^+/+^). Two uncoordinated adults visible in addition to sparse larvae retarded in development. Representative images captured reflecting the growth rate deficit (compare panel A and B; synchronized isogenic populations shown).

Since, the original lines expressing human ß-amyloid (Link 1995) appear only mildly affected in the temperature-sensitive manner, we further sought to compare the stable doubly-homozygous lines (expressing expanded polyQ-YFPs in addition to amyloid encoding *dvIs2*) looking for the temperature-dependent phenotypes indicative for aggravated proteotoxicity. As shown in Fig. 3 doubly-homozygous animals (*dvIs2* ^+/+;^ *rmIs133* ^+/+^) propagated at restrictive conditions develop temperature-dependent uncoordinated (*unc*) phenotype of a progressive palsy type and insidious onset. Presented results demonstrate the vestigial movement restricted to the animals head, persisting in otherwise uncoordinated individuals. Reported (Fig. 3A) temperature-sensitive effects are recollective of phenotypes presented previously (Link et al. 2003) in comparable model of acute (*smg-1ts* dependent) ß-amyloid proteotoxicity. We subsequently followed the temperature-sensitive effects we observed initially in isogenic populations of double-homozygotes, co-expressing ER-routed ß-amyloid and polyQ40-YFP propagated at 25°C for 10 days. In the laboratory settings isogenic *C. elegans* populations maintained on the solid agar media predate on the layer of *E. coli* cells spotted in the center of the plate. Evanescence of the bacterial feeding layer after several days of cultivation indicates for relatively unperturbed population growth. Conversely, the persistence of bacterial lawn is regarded as a vital parameter in genetic, RNAi, pathogen and xenobiotic assays. As presented in Fig. 3, we observed the persistence of the bacterial lawn in two concordant populations of doubly-homozygous (*dvIs2* ^+/+;^ *rmIs133* ^+/+^) genotype (derived from independent mating groups) at day 10 of growth at non-permissive temperature. We contrast the above outcome by comparing with any of the singly-transgenic homozygous parental lines [expressing either ß-amyloid (*dvIs2* ^+/+)^ or polyQ40-YFP (*rmIs133* ^+/+^), presented as insets in Fig. 3B and 3C] tested in parallel at 25°C. Above results suggest that, while the proteotoxic effects of ß-amyloid and expanded polyQ repeats are enhanced by the elevated temperatures, neither of the transgenes tested in singly-transgenic background is sufficient to account for severe growth decline observed when transgenes encoding for both pathogenic polypeptides are tested in combination.

**Figure 3.**
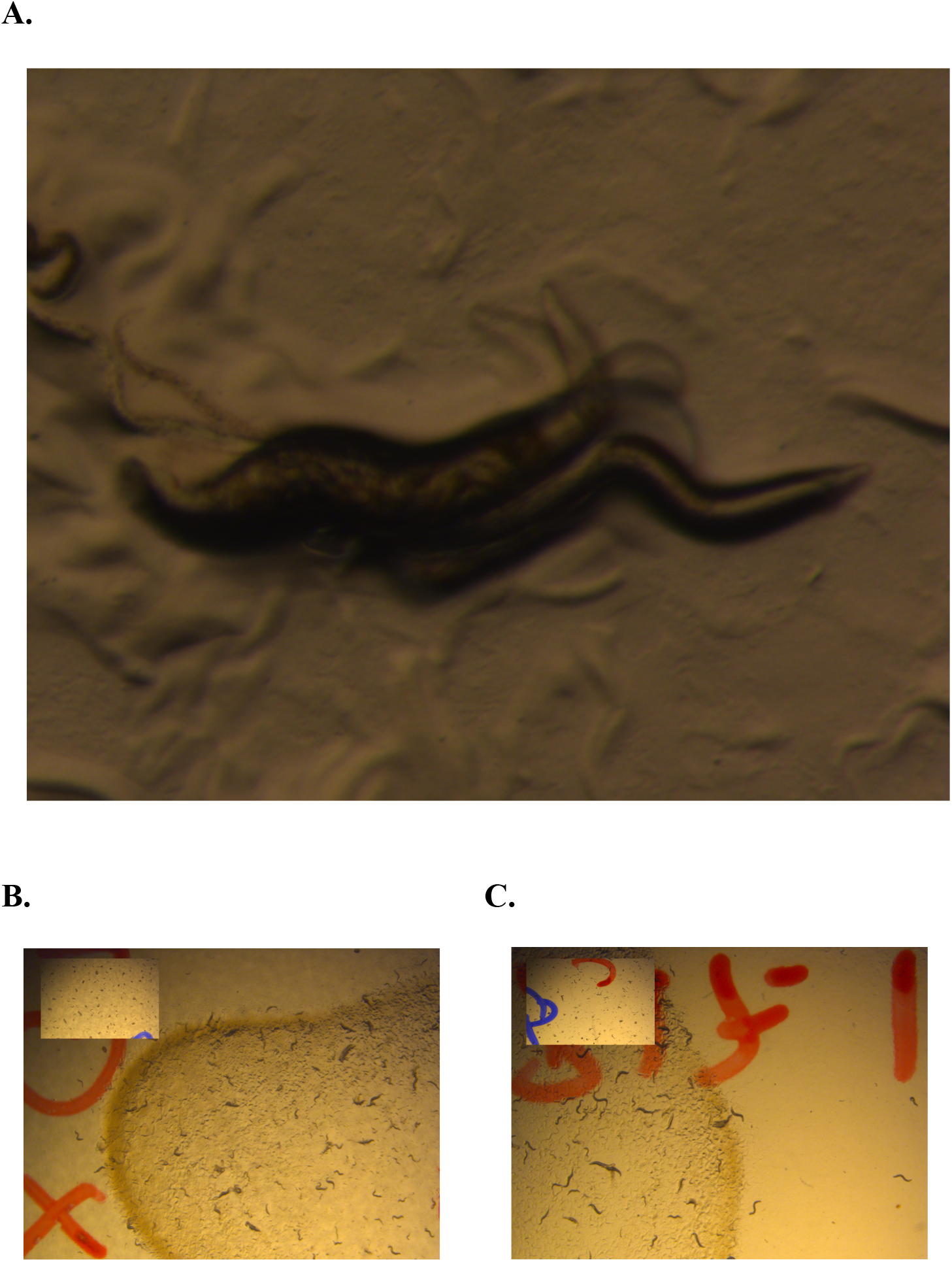
The uncoordinated phenotype of a progressive palsy type and insidious onset: temperature-sensitive effects of combined proteopathy consisting of transgenic ß-amyloid and expanded polyQ40-YFP (*dvIs2* ^+/+;^ *rmIs133* ^+/+^). A. Merged overlay of the sequential captures of the vestigial movement restricted to the animals head persisting in otherwise uncoordinated (immobile animal visible in the center) plegic adult. Compare to previously reported plate images (presented in Fig. 1D in Link et al. 2003). B and C. Plate phenotypes of admixed proteopathy recorded at 10 days of cultivation at 25°C. Apparent persistence of the bacterial lawn in two concordant populations of doubly-homozygous (*dvIs2* ^+/+;^ *rmIs133* ^+/+^) consistent with severe growth decline. Compare with appearance of the singly transgenic parental lines included as controls: ß-amyloid transgene (*dvIs2;* the inset on panel B) or polyQ40-YFP transgene (*rmIs133;* the inset on panel C). Note consistent saturation with starved larval stages on control plates presented as insets.

We recapitulated the observed temperature-sensitive phenotype of double-transgenic lines co-expressing ß-amyloid and polyQ40-YFP by means of RNAi mediated withdrawal of HSP70A (F26D10.3). Administration, via feeding route, of the dsRNA corresponding to *hsp-1* into the isogenic hermaphrodites of above double-transgenic background appeared to phenocopy the temperature-sensitive effect (data not shown, see discussion). We choose the HSP-1 since our previous report (Fonte et al. 2002) specified this particular homolog of human cytosolic HSP70 is one of the major human ß-amyloid interacting chaperone proteins in transgenic *C. elegans* expressing ER routed Aß42. In the descriptions of that work, we concluded, that in contrast to ER resident HSP70 chaperones *hsp-3 and hsp-4* (Kapulkin et al. 2005), cytosolic *hsp-1* is an essential gene strictly required for growth and development and therefore abstained from the in-depth characterization of subviable RNAi phenotypes. Unexpectedly, the attempts to reproduce those original results with combined proteopathy model failed – control animals (of singly transgenic parental genotypes expressing either ß-amyloid or polyQ-YFP) exposed to *hsp-1* RNAi were viable and superficially affected only mildly. The reasons possibly explaining the above inconsistency are stated and discussed in the following section.

Above phenotypes observed in combined proteopathy model prompted the following conclusion: polyQ40-YFP act synergistically with ß-amyloid and promotes temperature-dependent proteotoxicity in double-transgenic lines co-expressing both pathogenic variants confined to the same cellular type. Hence, conclusion tentatively consistent with cell-autonomous pathogenicity model. Therefore, we examined the subcellular distribution of ß-amyloid fibers with respect to inclusions formed by the aggregation-prone polyQ40-YFP. The results of confocal sectioning across sagittal plane of the (*dvIs2* ^+/+^; *rmIs133* ^+/+^) animal head (Fig. 4) demonstrate the cytosolic YFP labeled polyQ40 inclusions counter-stained *in vivo* with amyloid specific dye X-34 (Styren et al. 2000, Link et al. 2001; Ikonomovic et al. 2006). Apparent X-34 reactive congophilic intracellular mature ß-amyloid fibers (Fig. 4A. and D.) distinct from punctate cytosolic inclusions (Fig. 4B. and D.) labeled by YFP appended onto polyQ repeats expanded above the pathogenic threshold. Presented data establish both (*dvIs2* ^+/+^ and *rmIs133* ^+/+^) transgenes are efficiently expressed in the combined proteopathy model; excluding the dominant forms of ‘transgene incompatibilities’, such us transgenic silencing or interference within or between integrated transgenic arrays, could significantly contribute to histopathological phenotypes observed in doubly-homozygous background. Detected ER-rerouted ß-amyloid fibers fail to colocalize with punctate polyQ inclusions – an observation consistent with amyloid plaque formation and concomitant polyQ inclusion deposition underlying and pathognomic of the cellular degeneration.

**Figure 4.**
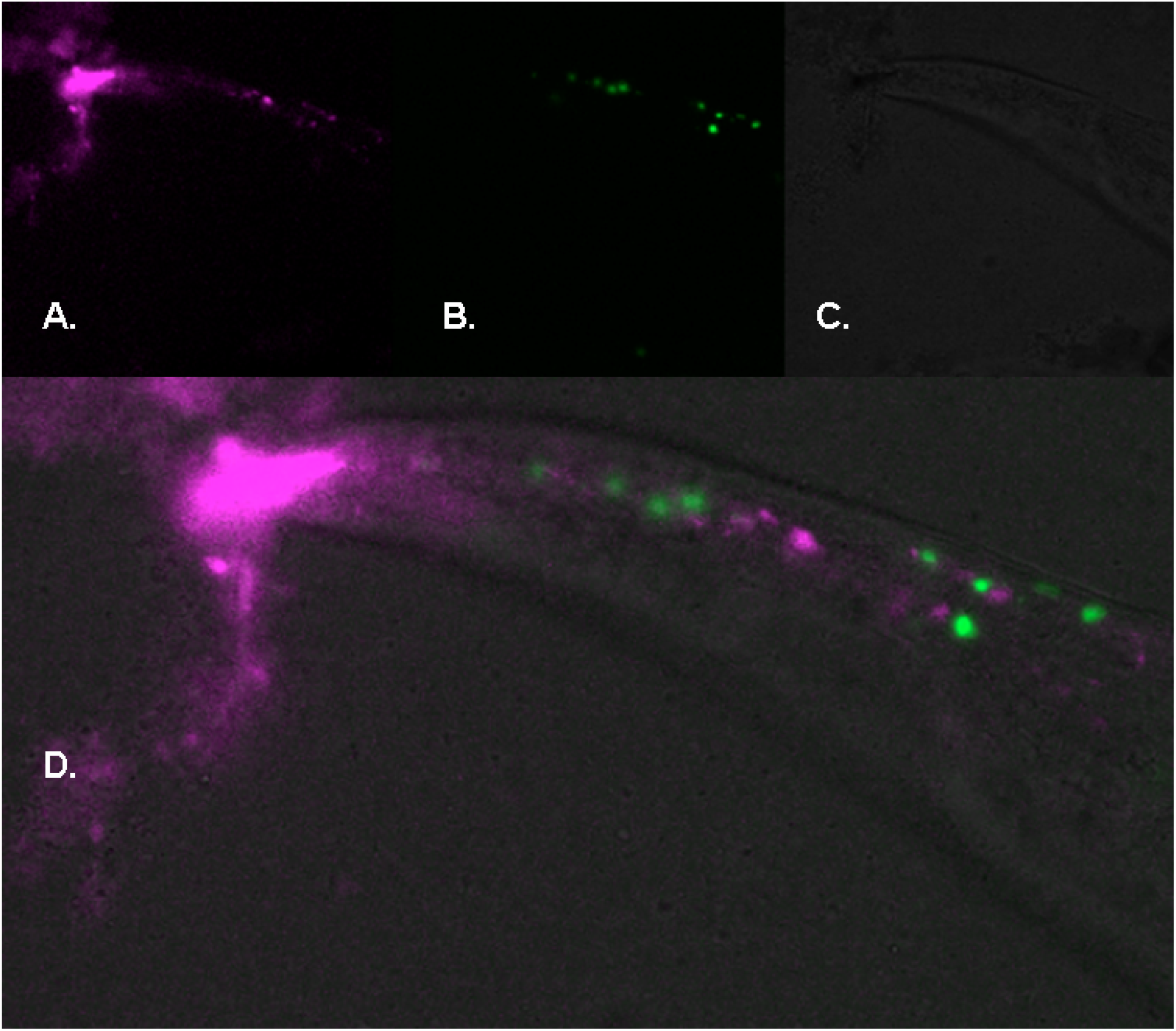
Histopathological phenotype of admixed proteopathy of Huntington and Alzheimer’s- type. Color-coded confocal section across sagittal plane of the animal head (8μm thick) expressing transgenic ß-amyloid and expanded polyQ40-YFP (*dvIs2* ^+/+;^ *rmIs133* ^+/+^). A. intracellular fibrillar ß-amyloid deposits distinctively reactive with X-34 (magenta). B. expanded polyQ40 appended with YFP (green). C. Brightfield contrast of the corresponding area. Congophilic intracellular mature ß-amyloid fibers (A. and D.) distinct from punctate cytosolic inclusions (B. and D.) formed by YFP appended onto polyQ repeats expanded above the pathogenic threshold. Detected ER- rerouted ß-amyloid fibers fail to colocalize with punctate polyQ inclusions – observation consistent with amyloid plaque formation and concomitant polyQ inclusion deposition underlying and pathognomic of the cellular degeneration. Note X-34 reactive mass situated in the rostral area composed of material attached to leftover undissolved stain.

Demonstrated ability to distinguish pathogenic polyQ inclusions from β-sheet secondary protein structures characteristic of X-34 staining in the living animal emphasizes the potential relevance of our combined proteopathy model. An important advantage of our model is the entire process of amyloid fibrillarization, plaque formation and polyQ deposition can be followed *in vivo*.

Results of confocal sectioning reveal X-34 reactive structures co-exist with morphologically distinct intracellular inclusions formed by the pathogenic polyQ repeats in a strictly cell-autonomous manner. This could be explained, provided the transgene encoding for human ß- amyloid produces ER-routed variant of β-peptide assumed to engage the intrinsic ER-cytoplasm retrograde transport (Fonte et al. 2002); mature fibers are observed as strictly intracellular structures found outside of the ER *i.e.* in the cytoplasm of cells affected by the combination of degenerative processes.

## DISCUSSION

We have created the *C. elegans* model for admixed Huntington’s and Alzheimer-type proteopathy. We found that transgenic mutant Huntington-type polyglutamine repeats expanded above pathogenic threshold co-exist with ß-amyloid containing senile plaques in a cell-autonomous manner. We established that the above disease associated aggregation-prone variants do form Huntington-type polyQ inclusions and congophilic Alzheimer’s-type fibrils confined to the cells expressing the *unc-54*. Strikingly our results revealed the admixed expression pattern consisting of apparently, non-overlapping YFP labeled polyglutamine aggregates and distinct X-34 reactive fibrillar ß-amyloid-type lesions. Provided the HD and AD-type proteinaceous foci can be distinguished by gross morphology and overall microscopic appearance *in vivo*, our results suggest that the aggregation of polyQ repeats and formation of mature ß-amyloid fibers may occur independently when the process is confined to the same cell. While, at the moment, we cannot entirely exclude that some smaller oligomeric variants might co-aggregate (either directly or by engaging interaction with common ligand – perceivably background events not captured at current resolution), we demonstrated that the prominent polyQ40-YFP aggregates occupy different sub-cellular territories and could be distinguished from mature ß-amyloid fibers.

Our results indicate that the co-occurrence of ß-amyloid fibers and extended polyglutamine Q40 aggregates invoked the synergistic lethality in a temperature-dependent manner. This means that in the model of combined proteopathy of AD and HD type, the two different aggregation-prone proteins may act cooperatively to promote the cell-autonomous proteotoxicity. While our results do not specify the direction of causality, the combined proteopathy model should be informative regarding the underlying pathological processes. Considering the pathways involved, we note the temperature sensitivity have been reported in *C. elegans* polyglutamine model (Gidalevitz et al. 2006). Their findings suggest that polyglutamine expansions impair the cellular protein folding capacity, hence affect the stability of temperature-sensitive mutant protein variants. Consistently aggregation of polyglutamines expanded above the pathogenic threshold (Morley et al. 2002) could aggravate the effects of ß-amyloid-type lesions as observed in our combined proteopathy model.

In that perspective expanded aggregation-prone polyglutamines coupled with nimbly folded fluorescent proteins could be regarded as ‘pathogenic probes’ useful in the proteotoxicity assays (*i.e.* involving ß-amyloid and possibly other comparable models of disease-associated pathogenic variants of aggregation-prone polypeptides), and provide the sensitized background for sophisticated screening. Combined pathogenic effects of different aggregation-prone polypetides could be considerred in terms of proteotoxic (-kinetic / -dynamic parameters) measures.’

Contemplating the screening strategies immediately available in our combined proteopathy model, we reported on the experimental setting based feasibility evaluation of the sensitized background screening approach utilizing the RNA interference (RNAi) - a popular screening mode applied for *C. elegans* genome (Fire et al. 1998). In particular, we observed, administration via feeding route, of dsRNA corresponding to a major ß-amyloid interacting protein *hsp-1* (Fonte et al. 2002), results in RNAi mediated silencing effect that recapitulate the temperature-sensitive proteotoxicity recorded in our combined proteopathy model. Verily others (Nollen et al. 2004), established the RNAi mediated withdrawal of *hsp-1* acts as a modulator of polyQ aggregation (Teuling et al. 2011). Remarkably, in a combinatorial screening for modifiers of ER-routed Aß42 effects Brehme et al. (2014) identified *hsp-1* RNAi among ATP-dependent chaperones that overlapped and enhanced proteotoxicity of both Aß42 and expanded polyglutamine transgenes. Results of RNAi screens concluded by Brehme et al. appear consistent with results of *hsp-1* withdrawal previously reported by Nollen et al. and align with results reported by Kraemer et al. (2006) specifying *hsp-1* RNAi as a modifier of pathogenic, aggregating human Tau variant expressed as transgene in *C. elegans* model for tauopathy. Conversely, independently carried RNAi screen reported by Khabirova et al. (2014) listed F26D10.3 (encoding for *hsp-1*) as a suppressor of Aß42 proteotoxicity observed in *C. elegans* AD model. Also, combinatorial chaperone HSP70 AMP-ylation study by Truttmann et al. (2018) indicated cytosolic *hsp-1* RNAi mediates the modulatory effects on aggregation and proteotoxicity of degenerative disease-associated polypeptides including ß-amyloid and polyQ. Could those miscellaneous and seemingly contradictory observations be reconciled in our combined proteopathy model? The effects of *hsp-1* RNAi withdrawal are expectedly pleiotropic and presumably cell-nonautonomous *i.e.* affect many tissues in addition to cells expressing proteotoxic polypeptides. Hence, some effects of *hsp-1* RNAi might be distant to tissue-specific response to abnormal protein aggregation. Additionally, given the complexity of the chaperone network, *hsp-1* RNAi withdrawal could affect the expression levels of other effector genes involved in the heat shock response – resulting in prominent secondary alternations of the compensatory type response as evidenced for ER-resident chaperones (Kapulkin et al. 2005). While RNAi mediated withdrawal of *hsp-1* does not significantly alter the ER resident *hsp-4* reporter (Calfon et al. 2002), RNAi against *hsp-1* does influence the levels of *hsp-16.2* reporter (Kapulkin WJ, Link CD 2002 unpublished observations; Guisbert et al. 2013)- an another effector chaperone we specified as directly interacting with human ß-amyloid (Fonte et al. 2002). Provided the accepted variability of the efficacy in experimental feeding of dsRNA silencing mediators we suggest one explanation would consider a dose-dependent model where the overall output of *hsp-1* RNAi withdrawal would perceivably involve alternations to gene expression at loci other than *hsp-1* accounting for secondary effects. Still, we note an alternative, yet not mutually exclusive, explanation might exist^1).^. Furthermore, we remark the above considerations concerning the results of RNAi-based screening could now be independently verified implementing the CRISPR/Cas9 improvement to generate precise variants (singly or in combination) in transiently silenced candidate genes including *hsp-1* discussed above. We have demonstrated previously that complex transgenic lines expressing Aß42 can be conveniently modified with CRISPR/Cas9 (Kapulkin et al. 2016), henceforth the stable transgenic lines reported here as the combined proteopathy model await further characterization.

Dementia is a historically recognized feature common in both HD and AD (Davis et al. 2014). Dementias are often classified as cortical (AD) or subcortical (HD) with symptoms reflecting the areas of the brain affected by degenerative lesions resulting in synaptic and neuronal loss (Salmon et al. 2007) leading to eventual neural system failure which drives the progression of the symptoms. The occurrence of degenerative lesions in HD and AD is a common feature in both conditions (Spires &Hannan 2007) however, due to very different aetiology and underlying genetics - the exact sites of histopathologically detected proteinaceous deposits overlap only partially, depending on the disease stage and a grade (Vonsatel et al. 1985; Braak&Braak 1991). Provided in HD and AD the degenerative process impairs the specific neuronal subtypes – distinction relevant particularly in prodromal and incipient disease progression stages - we designed an experimental settings and utilized a versatile genetic model system to combine both Huntington and Alzheimer-type lesions in cell-autonomous manner. Our two-component model combines the established Aß42-dependent amyloid root of an AD (Selkoe 1997; Hamley 2012; Jonsson et al. 2012) and expanded polyglutamine root of HD (Walker 2007; Finkbeiner 2011). Based on the combination of two convergent degenerative pathways, we concluded polyglutamines expanded above pathogenic threshold exacerbates the Aß42-amyloid effects observable in our model; however, we do not know the direction of causality neither we do fully understand the pathological process underlying the disease mechanism. While presented results might suffer from insufficient characterization in terms of quantitative measures applied (to strictly quantify the observed phenotypes e.g. uncoordinated movement, aggregates number etc), we provided the evidence for synchronous expression of ß- amyloid and expanded polyglutamine reporter consistent with characterized *unc-54* promoter specificity.

Combined proteopathies have been described in man (McIntosh et al. 1978; Vonsattel et al. 1985; Reyes &Gibbons 1985; Moss et al. 1988; Bruyn &Roos 1990; Becher et al. 1997; and Jellinger 1998 with relevants references therein), however, histopathologically confirmed descriptions of Alzheimer-type lesions in HD are sparse (Davis et al. 2014; St-Amour et al. 2017). We note, our model could be directly informative for those admixed proteopathies. Further, we perceive the cell-autonomous readout might define the common denominator for the underlying disease mechanisms considered singly. Histopathologically HD and AD share extensive parallels observed at the cellular level (reviewed in Spires et al. 2007) albeit of different aetiology and underlying genetics. Despite the major socio-economic impact, the pathogenesis of those diseases is not understood sufficiently neither the available therapeutic interventions are efficient.

We have previously established that *C. elegans* ß-amyloid model recapitulate the key Alzheimer-type chaperone response (Fonte et al. 2002), while others have demonstrated the induction of chaperone response in *C. elegans* model for expanded polyglutamines (Satyal et al. 2000) - apparently a convergent pathogenic feature recognized in AD and HD brain (Ehrnhoefer et al. 2011; Wyttenbach &Arrigo 2013; Brehme &Voisine 2016). Chaperones may act as promiscuous ligands, binding both expanded polyglutamines and Aß42 in oligomeric forms, hence our combined proteopathy model might provide a better controlled background for future studies (as previously Fonte et al. utilized an unlocalized GFP as a co-IP specificity control). Moreover, our previous report implicated the retrograde transport of ER-routed Aß42 is responsible for cell-autonomous ß- amyloid formation. This appears possibly relevant, since certain perturbations to ER proteostasis have a protective effect against ß-amyloid toxicity in *C. elegans* AD model (Kapulkin et al 2005; Lopez 2009). Protective effects against polyglutamine toxicity, involving the modulation of chaperone protein response, have been reported in other model systems (reviewed in Brehme &Voisine 2016). Together, we perceive the chaperone type response is invoked in our combined proteopathy model, however the full spectrum of the response awaits further characterization. Collectively the data argue in favor of the temperature-sensitive dose-dependent proteotoxicity model involving the depletion of the chaperone system capacity required to buffer the combined load of the aggregation-prone cargo. Given the preliminary nature of our report, presently we can not conclusively exclude the other pathogenesis mediators, such us postulated RNA-repeat toxicity pathways (Nalavade et al. 2013; Garcia et al. 2014; Marti 2016), imposes on and contributes to the overall combined proteopathy phenotype (Li et al. 2008). We note however, that overall length of expanded polyQ encoding (*CAG)* repeats used in our model might not suffice for significant RNA-repeat toxicity (Finkbeiner 2011).

A different type of combined proteopathy – of co-occurring Huntington and Parkinson type - have been described in a fly model (Pocas et al. 2015). That model combines the effects of α- synuclein on aggregation and toxicity of expanded polyglutamines. In that model α-synuclein and polyQ do form co-aggregates and mutually modulate the proteotoxic effects. While our findings are in a substantial agreement with proposed synergistic (or additive) actions of both aggregation-prone proteins *in vivo*, the co-aggregation of α-synuclein and polyQ is contrasted with aggregative behaviour of extended polyglutamines in the presence of amyloid-plaques observed in our model. Our results are consistent with the co-existence of two distinct types of proteinaceous foci; supporting the perspective when polyglutamines appended with fluorescent protein form one type of inclusion bodies (composed of less-ordered nonamyloid protein aggregates), and the other type of lesions forming X-34 reactive mature amyloid fibrils. While insoluble fibril-like structures have been attributed to aggregating polyglutamines (Sipe et al. 2016; OMIM#143100 and references therein) those ultrastructural characteristics do not confer properties of AD-type congophilic fibrills (DiFiglia et al. 1997; OMIM#104300 and references therein). As reviewed by Finkbeiner (2011) expanded polyglutamines in HD result in the visible protein accumulations, but the Huntington-type inclusion bodies are composed mostly of less-ordered nonamyloid protein aggregates.

Collectively, the main finding inferred from our combined proteopathy model suggests the Huntington-type polyQ inclusions are composed of substantially amorphous granular material distinct from mostly fibrillar AD-type X-34 reactive amyloid-plaques. Although X-34 stains *post-mortem* human AD brains, showing 1:1 correspondence of X-34 and anti-Aß antibody staining of plaques and cerebrovascular amyloid (Styren et al. 2000), our *in vivo* cell-autonomous model of combined proteopathy revealed an additional layer of proteotoxic complexity. Due to advantages of the cell-autonomous model, we uncovered at least two different types of intracellular inclusions: amyloidic and non-amyloidic may co-exists *in vivo*, supporting the two-state aggregation model of pathogenesis. We suggest the accumulation of aberrant polyQ containing proteins need to be taken into account as possibly contributing factor in AD, but also in other disorders associated with increased Aß aggregation and senile plaque formation. Most notably neurodegenerative progression in Down syndrome (OMIM#190685) may be affected by perceivably rare concomitant expansion of non-amyloidic polyglutamine stretches.

Lastly, we note, our results corroborate with and pertain to several lines of available literature evidence: *i.*) apparently pathogenic, non-amyloidic, non-fibrillar, non-congophilic Aß plaques were detected in *C. elegans* (Link et al. 2003), *Drosophila* (Crowther et al. 2005) AD models, and were described in certain cases of AD (Kumar-Singh et al. 2000) linked to APP variant (T714I). *ii.*) Singhrao et al. (1998) established that in AD immuno-histochemistry with anti-HTT antibody localized only with the intracellular structures but not with the mature amyloid. *iii.*) heritable polyQ disorders other than Huntington’s chorea were proposed as contributing factor in AD (Reid et al. 2004) and TDP-43 dependent ALS (Elden et al. 2010).

## Acknowledgments

We are grateful to Mihail Sarov and Anthony Hyman for hosting us at MPI-CBG. We are grateful to William E. Klunk (University of Pittsburgh Department of Psychiatry – UPMC) for providing us with X-34 stain. We are grateful to Richard Morimoto (Northwestern University) and members of the Morimoto laboratory (Susan Fox and Renee Brielmann) for providing the polyQ-YFP strains and constructs. We are grateful to Paul Francis (Institute of Psychiatry, Psychology & Neuroscience, King’s College London) and Kurt Jellinger (Institute of Clinical Neurobiology, Vienna) for insightful discussions concerning the combined proteinopathies. We are grateful to Andrew Fire (Stanford) for discussions concerning RNA interference. We acknowledge Jan Peychl (MPI-CBG Light Microscopy Facility) for expertise in confocal microscopy. We note, some strains were provided by the CGC, which is funded by NIH Office of Research Infrastructure Programs (P40 OD010440), and the RNAi strains and constructs were provided by Silencing Genomes resource at the Cold Spring Harbor Laboratory.

## Footnotes

1). While we can not exclude that the variability in the strength of RNAi could be affected by procedural differences (affecting the rate of cellular synthesis of siRNA, factors affecting the dsRNA uptake or stability, levels of T7 polymerase induction etc), we note the particular dsRNA encoding plasmids used were designed differently. The original report (Fonte et al. 2002) utilized constructed plasmids entailing the entire gene coding segment of *hsp-1* including introns. Instead the later efforts relied on the construct distributed with commonly available RNAi feeding library (Kamath et al. 2003) incorporating considerably smaller fragment of *hsp-1* gene.

## BIBLIOGRAPHY

1. Beam, M, Silva, MC, Morimoto, RI. Dynamic imaging by fluorescence correlation spectroscopy identifies diverse populations of polyglutamine oligomers formed in vivo. J Biol Chem. 2012 Jul 27;287(31):26136–45.

2. Becher MW, Kolzuk JA, Wagster MV et al. 1997 Amyloid deposits and neuritic plaques in Huntington’s disease. Journal of Neuropathology and Experimental Neurology 56(5)

3. Braak H, Braak E. Neuropathological stageing of Alzheimer-related changes. Acta Neuropathol. 1991;82(4):239–59. Review. PubMed PMID: 1759558.

4. Brehme M, Voisine C, Rolland T, Wachi S, Soper JH, Zhu Y, Orton K, Villella A, Garza D, Vidal M, Ge H, Morimoto RI. A chaperome subnetwork safeguards proteostasis in aging and neurodegenerative disease. Cell Rep. 2014 Nov 6;9(3):1135–50. doi: 10.1016/j.celrep.2014.09.042. Epub 2014 Oct 23. PubMed PMID: 25437566; PubMed Central PMCID: PMC4255334.

5. Brehme M, Voisine C. Model systems of protein-misfolding diseases reveal chaperone modifiers of proteotoxicity. Dis Model Mech. 2016 Aug 1;9(8):823–38. doi: 10.1242/dmm.024703. Review. PubMed PMID: 27491084; PubMed Central PMCID: PMC5007983.

6. Brenner S. The genetics of Caenorhabditis elegans. Genetics. 1974 May;77(1):71–94.

7. Brignull, H.R., J.F. Morley, S.M. Garcia, and R.I. Morimoto. Modeling Polyglutamine Pathogenesis in C. elegans. Methods in Enzymology 412: 256–282 (2006).

8. Bruyn GW, Roos RA. Senile plaques in Huntington’s disease: a preliminary report. Clin Neurol Neurosurg. 1990;92(4):329–31. PubMed PMID: 1963823.

9. Calfon M, Zeng H, Urano F, Till JH, Hubbard SR, Harding HP, Clark SG, Ron D. IRE1 couples endoplasmic reticulum load to secretory capacity by processing the XBP-1 mRNA. Nature. 2002 Jan 3;415(6867):92–6. Erratum in: Nature 2002 Nov 14;420(6912):202. PubMed PMID: 11780124.

10. Carrell RW, Lomas DA. Conformational disease. Lancet. 1997 Jul 12;350(9071):134–8. Review. PubMed PMID: 9228977.

11. Crowther DC, Kinghorn KJ, Miranda E, Page R, Curry JA, Duthie FA, Gubb DC, Lomas DA. Intraneuronal Abeta, non-amyloid aggregates and neurodegeneration in a Drosophila model of Alzheimer’s disease. Neuroscience. 2005;132(1):123–35. PubMed PMID: 15780472.

12. Davis MY, Keene CD, Jayadev S, Bird T. The co-occurrence of Alzheimer’s disease and Huntington’s disease: a neuropathological study of 15 elderly Huntington’s disease subjects. J Huntingtons Dis. 2014;3(2):209–17. doi: 10.3233/JHD-140111. PubMed PMID: 25062863.

13. DiFiglia M, Sapp E, Chase KO, Davies SW, Bates GP, Vonsattel JP, Aronin N. Aggregation of huntingtin in neuronal intranuclear inclusions and dystrophic neurites in brain. Science. 1997 Sep 26;277(5334):1990–3. PubMed PMID: 9302293.

14. Ehrnhoefer, D. E., Wong, B. K., & Hayden, M. R. (2011). Convergent pathogenic pathways in Alzheimer’s and Huntington’s diseases: shared targets for drug development. Nature reviews. Drug discovery, 10(11), 853–867. doi:10.1038/nrd3556

15. Elden AC, Kim HJ, Hart MP, Chen-Plotkin AS, Johnson BS, Fang X, Armakola M, Geser F, Greene R, Lu MM, Padmanabhan A, Clay-Falcone D, McCluskey L, Elman L, Juhr D, Gruber PJ, Rüb U, Auburger G, Trojanowski JQ, Lee VM, Van Deerlin VM, Bonini NM, Gitler AD. Ataxin-2 intermediate-length polyglutamine expansions are associated with increased risk for ALS. Nature. 2010 Aug 26;466(7310):1069–75. doi: 10.1038/nature09320. PubMed PMID: 20740007; PubMed Central PMCID: PMC2965417.

16. Finkbeiner S. Huntington’s Disease. Cold Spring Harb Perspect Biol. 2011 Jun 1;3(6). pii: a007476. doi: 10.1101/cshperspect.a007476. Review. PubMed PMID: 21441583; PubMed Central PMCID: PMC3098678.

17. Fire A, Xu S, Montgomery MK, Kostas SA, Driver SE, Mello CC. Potent and specific genetic interference by double-stranded RNA in Caenorhabditis elegans. Nature. 1998 Feb 19;391(6669):806–11. PubMed PMID: 9486653.

18. Fonte V, Kapulkin WJ, Taft A, Fluet A, Friedman D, Link CD. Interaction of intracellular beta amyloid peptide with chaperone proteins. Proc Natl Acad Sci U S A. 2002 Jul 9;99(14):9439–44. Epub 2002 Jun 27. Erratum in: Proc Natl Acad Sci U S A. 2013 Mar 19;110(12):4853.

19. Garcia SM, Tabach Y, Lourenço GF, Armakola M, Ruvkun G. Identification of genes in toxicity pathways of trinucleotide-repeat RNA in C. elegans. Nat Struct Mol Biol. 2014 Aug;21(8):712–20. doi: 10.1038/nsmb.2858. Epub 2014 Jul 20. PubMed PMID: 25038802; PubMed Central PMCID: PMC4125460.

20. Gidalevitz T, Ben-Zvi A, Ho KH, Brignull HR, Morimoto RI. Progressive disruption of cellular protein folding in models of polyglutamine diseases. Science. 2006 Mar 10;311(5766):1471–4. Epub 2006 Feb 9. PubMed PMID: 16469881.

21. Guisbert E, Czyz DM, Richter K, McMullen PD, Morimoto RI. Identification of a tissue-selective heat shock response regulatory network. PLoS Genet. 2013 Apr;9(4):e1003466. doi: 10.1371/journal.pgen.1003466. Epub 2013 Apr 18. PubMed PMID: 23637632; PubMed Central PMCID: PMC3630107.

22. Hamley IW. The amyloid beta peptide: a chemist’s perspective. Role in Alzheimer’s and fibrillization. Chem Rev. 2012 Oct 10;112(10):5147–92. doi: 10.1021/cr3000994. Epub 2012 Jul 19. Review. PubMed PMID: 22813427.

23. Hodgkin J, Horvitz HR, Brenner S. Nondisjunction Mutants of the Nematode CAENORHABDITIS ELEGANS. Genetics. 1979 Jan;91(1):67–94.

24. Ikonomovic MD, Abrahamson EE, Isanski BA, Debnath ML, Mathis CA, Dekosky ST, Klunk WE. X-34 labeling of abnormal protein aggregates during the progression of Alzheimer’s disease. Methods Enzymol. 2006;412:123–44. Review. PubMed PMID:17046656.

25. Jellinger KA. Alzheimer-type lesions in Huntington’s disease. J Neural Transm (Vienna). 1998;105(8-9):787–99. PubMed PMID: 9869319.

26. Jonsson T, Atwal JK, Steinberg S, Snaedal J, Jonsson PV, Bjornsson S, Stefansson H, Sulem P, Gudbjartsson D, Maloney J, Hoyte K, Gustafson A, Liu Y, Lu Y, Bhangale T, Graham RR, Huttenlocher J, Bjornsdottir G, Andreassen OA, Jönsson EG, Palotie A, Behrens TW, Magnusson OT, Kong A, Thorsteinsdottir U, Watts RJ, Stefansson K. A mutation in APP protects against Alzheimer’s disease and age-related cognitive decline. Nature. 2012 Aug 2;488(7409):96–9. doi: 10.1038/nature11283. PubMed PMID: 22801501.

27. Kamath RS, Ahringer J. Genome-wide RNAi screening in Caenorhabditis elegans. Methods. 2003 Aug;30(4):313–21. PubMed PMID: 12828945.

28. Kapulkin WJ, Hiester BG, Link CD. Compensatory regulation among ER chaperones in C. elegans. FEBS Lett. 2005 Jun 6;579(14):3063–8. Erratum in: FEBS Lett. 2007 Dec 22;581(30):5952

29. Kapulkin W, Sarov M. Allelic spectrum of the RNA guided CRISPR/Cas9 DNA repair events at PAM associated trinucleotide repeat (NGG)n in Caenorhabditis elegans. bioRxiv 062489; 10 Jul 2016. doi: https://doi.org/10.1101/062489

30. Khabirova E, Moloney A, Marciniak SJ, Williams J, Lomas DA, Oliver SG, Favrin G, Sattelle DB, Crowther DC. The TRiC/CCT chaperone is implicated in Alzheimer’s disease based on patient GWAS and an RNAi screen in Aβ-expressing Caenorhabditis elegans. PLoS One. 2014 Jul 31;9(7):e102985. doi: 10.1371/journal.pone.0102985. eCollection 2014. PubMed PMID: 25080104; PubMed Central PMCID: PMC4117641.

31. Kopito RR, Ron D. Conformational disease. Nat Cell Biol. 2000 Nov;2(11):E207–9. PubMed PMID: 11056553.

32. Kraemer BC, Burgess JK, Chen JH, Thomas JH, Schellenberg GD. Molecular pathways that influence human tau-induced pathology in Caenorhabditis elegans. Hum Mol Genet. 2006 May 1;15(9):1483–96. Epub 2006 Apr 6. PubMed PMID: 16600994.

33. Kumar-Singh S, De Jonghe C, Cruts M, Kleinert R, Wang R, Mercken M, De Strooper B, Vanderstichele H, Löfgren A, Vanderhoeven I, Backhovens H, Vanmechelen E, Kroisel PM, Van Broeckhoven C. Nonfibrillar diffuse amyloid deposition due to a gamma(42)-secretase site mutation points to an essential role for N-truncated A beta(42) in Alzheimer’s disease. Hum Mol Genet. 2000 Nov 1;9(18):2589–98. PubMed PMID: 11063718.

34. Li LB, Yu Z, Teng X, Bonini NM. RNA toxicity is a component of ataxin-3 degeneration in Drosophila. Nature. 2008 Jun 19;453(7198):1107–11. doi:10.1038/nature06909. Epub 2008 Apr 30. PubMed PMID: 18449188; PubMed Central PMCID: PMC2574630.

35. Link CD. Expression of human beta-amyloid peptide in transgenic Caenorhabditis elegans. Proc Natl Acad Sci U S A. 1995 Sep 26;92(20):9368–72. PubMed PMID: 7568134; PubMed Central PMCID: PMC40986.

36. Link CD, Johnson CJ, Fonte V, Paupard M, Hall DH, Styren S, Mathis CA, Klunk WE. Visualization of fibrillar amyloid deposits in living, transgenic Caenorhabditis elegans animals using the sensitive amyloid dye, X-34. Neurobiol Aging. 2001 Mar-Apr;22(2):217–26. PubMed PMID: 11182471.

37. Link CD, Taft A, Kapulkin WJ, Duke K, Kim S, Fei Q, Wood DE, Sahagan BG. Gene expression analysis in a transgenic Caenorhabditis elegans Alzheimer’s disease model. Neurobiol Aging. 2003 MayJun;24(3):397–413. [correction pending]

38. Lopez, Nunez Yury Orlando (2009). UDP-glucose glycoprotein glucosyltransferase (uggt-1) and UPR genes modulate C. elegans necrotic cell death. Dissertation under Prof. Monica Driscoll PhD. Rutgers University.

39. Martí E. RNA toxicity induced by expanded CAG repeats in Huntington’s disease. Brain Pathol. 2016 Nov;26(6):779–786. doi: 10.1111/bpa.12427. Review. PubMed PMID: 27529325.

40. McIntosh GC, Jameson HD, Markesbery WR. Huntington disease associated with Alzheimer disease. Ann Neurol. 1978 Jun;3(6):545–8. PubMed PMID: 150253.

41. Moss RJ, Mastri AR, Schut LJ. The coexistence and differentiation of late onset Huntington’s disease and Alzheimer’s disease. A case report and review of the literature. Journal of the American Geriatrics Society [01 Mar 1988, 36(3):237–241]

42. Morley JF, Brignull HR, Weyers JJ, Morimoto RI. The threshold for polyglutamine expansion protein aggregation and cellular toxicity is dynamic and influenced by aging in Caenorhabditis elegans. Proc Natl Acad Sci U S A 99: 10417–22, 2002.

43. Nalavade R, Griesche N, Ryan DP, Hildebrand S, Krauss S. Mechanisms of RNA-induced toxicity in CAG repeat disorders. Cell Death Dis. 2013 Aug 1;4(8):e752. doi: 10.1038/cddis.2013.276. PubMed PMID: 23907466; PubMed Central PMCID: PMC3763438.

44. Nollen EA, Garcia SM, van Haaften G, Kim S, Chavez A, Morimoto RI, Plasterk RH. Genome-wide RNA interference screen identifies previously undescribed regulators of polyglutamine aggregation. Proc Natl Acad Sci U S A. 2004 Apr 27;101(17):6403–8. Epub 2004 Apr 14. PubMed PMID: 15084750; PubMed Central PMCID: PMC404057.

45. Poças GM, Branco-Santos J, Herrera F, Outeiro TF, Domingos PM. α-Synuclein modifies mutant huntingtin aggregation and neurotoxicity in Drosophila. Hum Mol Genet. 2015 Apr 1;24(7):1898–907. doi: 10.1093/hmg/ddu606. Epub 2014 Dec 1. PubMed PMID: 25452431; PubMed Central PMCID: PMC4355023.

46. Reid SJ, van Roon-Mom WM, Wood PC, Rees MI, Owen MJ, Faull RL, Dragunow M, Snell RG. TBP, a polyglutamine tract containing protein, accumulates in Alzheimer’s disease. Brain Res Mol Brain Res. 2004 Jun 18;125(1-2):120–8. PubMed PMID: 15193429.

47. Reyes MG, Gibbons S. Dementia of the Alzheimer’s type and Huntington’s disease. Neurology. 1985 Feb;35(2):273–7. PubMed PMID: 3155827.

48. Ron D. Proteotoxicity in the endoplasmic reticulum: lessons from the Akita diabetic mouse. J Clin Invest. 2002 Feb;109(4):443–5. Review. PubMed PMID: 11854314; PubMed Central PMCID: PMC150880.

49. Salmon DP, Filoteo JV. Neuropsychology of cortical versus subcortical dementia syndromes. Semin Neurol. 2007 Feb;27(1):7–21. Review. PubMed PMID: 17226737.

50. Satyal SH, Schmidt E, Kitagawa K, Sondheimer N, Lindquist S, Kramer JM, Morimoto RI. Polyglutamine aggregates alter protein folding homeostasis in Caenorhabditis elegans. Proc Natl Acad Sci U S A. 2000 May 23;97(11):5750–5. PubMed PMID: 10811890; PubMed Central PMCID: PMC18505.

51. Schindelin, J.; Arganda-Carreras, I. & Frise, E. et al. (2012), “Fiji: an open-source platform for biological-image analysis”, Nature methods 9(7): 676–682, PMID 22743772, doi:10.1038/nmeth.2019

52. Selkoe DJ. Alzheimer’s disease: genotypes, phenotypes, and treatments. Science. 1997 Jan 31;275(5300):630–1. Review. PubMed PMID: 9019820.

53. Singhrao SK, Thomas P, Wood JD, MacMillan JC, Neal JW, Harper PS, Jones AL. Huntingtin protein colocalizes with lesions of neurodegenerative diseases: An investigation in Huntington’s, Alzheimer’s, and Pick’s diseases. Exp Neurol. 1998 Apr;150(2):213–22. PubMed PMID: 9527890.

54. Sipe JD, Benson MD, Buxbaum JN, Ikeda SI, Merlini G, Saraiva MJ, Westermark P. Amyloid fibril proteins and amyloidosis: chemical identification and clinical classification International Society of Amyloidosis 2016 Nomenclature Guidelines. Amyloid. 2016 Dec;23(4):209–213. Epub 2016 Nov 24. PubMed PMID: 27884064.

55. Spires TL, Hannan AJ. Molecular mechanisms mediating pathological plasticity in Huntington’s disease and Alzheimer’s disease. J Neurochem. 2007 Feb;100(4):874–82. Epub 2007 Jan 8. Review. PubMed PMID: 17217424.

56. St-Amour I, Turgeon A, Goupil C, Planel E, Hébert SS. Co-occurrence of mixed proteinopathies in late-stage Huntington’s disease. Acta Neuropathol. 2018 Feb;135(2):249–265. doi: 10.1007/s00401-017-1786-7. Epub 2017 Nov 13. PubMed PMID: 29134321.

57. Stroo E, Koopman M, Nollen EA and Mata-Cabana A (2017) Cellular Regulation of Amyloid Formation in Aging and Disease. Front. Neurosci. 11:64. doi: 10.3389/fnins.2017.00064

58. Styren SD, Hamilton RL, Styren GC, Klunk WE. X-34, a fluorescent derivative of Congo red: a novel histochemical stain for Alzheimer’s disease pathology. J Histochem Cytochem. 2000 Sep;48(9):1223–32. PubMed PMID: 10950879.

59. Tanskanen M, “Amyloid” — Historical Aspects. In: Dali Feng (Eds.), Amyloidosis, 2013 https://doi.org/10.5772/53423

60. Teuling E, Bourgonje A, Veenje S, Thijssen K, de Boer J, van der Velde J, Swertz M, Nollen E. Modifiers of mutant huntingtin aggregation: functional conservation of C. elegans-modifiers of polyglutamine aggregation. PLoS Curr. 2011 Aug 12;3:RRN1255. doi: 10.1371/currents.RRN1255. PubMed PMID: 21915392; PubMed Central PMCID: PMC3166184.

61. Timmons L, Court DL, Fire A. Ingestion of bacterially expressed dsRNAs can produce specific and potent genetic interference in Caenorhabditis elegans. Gene. 2001 Jan 24;263(1-2):103–12. PubMed PMID: 11223248.

62. Truttmann MC, Pincus D, Ploegh HL. Chaperone AMPylation modulates aggregation and toxicity of neurodegenerative disease-associated polypeptides. Proc Natl Acad Sci U S A. 2018 May 29;115(22):E5008–E5017. doi: 10.1073/pnas.1801989115. Epub 2018 May 14. PubMed PMID: 29760078; PubMed Central PMCID: PMC5984528.

63. Vonsattel JP, DiFiglia M. Huntington disease. J Neuropathol Exp Neurol. 1998 May;57(5):369–84. Review. PubMed PMID: 9596408.

64. Vonsattel JP, Myers RH, Stevens TJ, Ferrante RJ, Bird ED, Richardson EP Jr. Neuropathological classification of Huntington’s disease. J Neuropathol Exp Neurol. 1985 Nov;44(6):559–77. PubMed PMID: 2932539.

65. Walker FO. Huntington’s disease. Lancet. 2007 Jan 20;369(9557):218–28. Review. PubMed PMID: 17240289.

66. Warner J, Barron L, St Clair D, Brock D. Reliability of clinical diagnosis of Huntington’s disease. J Neurol Neurosurg Psychiatry. 1994 Oct;57(10):1277. PubMed PMID: 7931402; PubMed Central PMCID: PMC485509.

67. Wenk GL. Neuropathologic changes in Alzheimer’s disease. J Clin Psychiatry. 2003;64 Suppl 9:7–10. Review. PubMed PMID: 12934968.

68. Wood WB (Eds.) 1988, Volume 17; The Nematode Caenorhabditis elegans. (CSHL monograph) Methods (Appendix) J. Sulston, J. Hodgkin

69. Wyttenbach A, Arrigo AP. The Role of Heat Shock Proteins during Neurodegeneration in Alzheimer’s, Parkinson’s and Huntington’s Disease. In: Madame Curie Bioscience Database [Internet]. Austin (TX): Landes Bioscience; 2000-2013. https://www.ncbi.nlm.nih.gov/books/NBK6495/

